# Aird: A computation-oriented mass spectrometry data format enables higher compression ratio and less decoding time

**DOI:** 10.1101/2020.10.14.338921

**Authors:** Miaoshan Lu, Shaowei An, Ruimin Wang, Jinyin Wang, Changbin Yu

## Abstract

With the precision of mass spectrometer going higher and the emergence of data independence acquisition (DIA), the file size is increasing rapidly. Beyond the widely-used open format mzML (Deutsch 2008), near-lossless or lossless compression algorithms and formats have emerged. The data precision is often related to the instrument and subsequent processing algorithms. Unlike storage-oriented formats, which focusing more on lossless compression and compression rate, computation-oriented formats focus as much on decoding speed and disk read strategy as compression rate. Here we describe “Aird", an opensource and computation-oriented format with controllable precision, flexible indexing strategies and high compression rate. Aird uses JavaScript Object Notation (JSON) for metadata storage, multiple indexing, and reordered storage strategies for higher speed of data randomly reading. Aird also provides a novel compressor called Zlib-Diff-PforDelta (ZDPD) for m/z data compression. Compared with Zlib only, m/z data size is about 65% lower in Aird, and merely takes 33% decoding time.

**Availability:** Aird SDK is written in Java, which allow scholars to access mass spectrometry data efficiently. It is available at https://github.com/Propro-Studio/Aird-SDK AirdPro can convert vendor files into Aird files, which is available at https://github.com/Propro-Studio/AirdPro

## Introduction

As integrity biological digital samples, vendor files are ideal for long-term storage for their high compression rate. However, due to cross-platform compatibility and software adaptation differences, Converting vendor files to other formats before data analysis is necessary. It is often discussed that the accuracy of converted files is lossless or near-lossless. Data precision is mainly determined by the accuracy of the mass spectrometer and analytical parameters, rather than the data digits stored. Using too many digits for calculation leads to a waste of computing resources, as well as lower calculation speed and software instability. Large files also bring a high cost of memory and bandwidth. In recent research, there are two directions on mass spectrometry (MS) data compression. One is the development of a new file format, the other is an exploration of a better data compressor. The MS-Numpress(Teleman, Dowsey et al. 2014) is a set of compressors providing algorithms with controllable precision. Considering the difference in the precision of intensity and m/z, MS-Numpress provides different near-lossless compression strategies. MassComp (Yang, Chen et al. 2019) presents a lossless compressor target on the m/z dimension, and it only works on mzXML (Pedrioli, Eng et al. 2004). The above two approaches are based on the mzXML or mzML(Deutsch 2008) format, which is unable to change the index of the spectrum data. Mz5(Wilhelm, Kirchner et al. 2012) and Toffee(Tully 2020) are two new file formats. They both use HDF5 as their core storage technology. MzDB(Bouyssie, Dubois et al. 2015) uses SQLite database technology for storage, which makes the random access file reading more flexible and simple. However, without specific libraries or software, these files are not readable (Teleman, Dowsey et al. 2014). A new format often brings much more additional work, such as adaptation to existing analysis software, build-up of software for file conversion, and compatibility with existing controllable vocabularies. However, when we use these data for calculation rather than long-term preservation, it is necessary to provide data with compatible precision and metadata information for software. Data not related to analysis is removed so as to reduce the cost of bandwidth and memory. In addition, the developers of the analysis software and the new format are often not the same. Therefore, the update progress of them are always not in accordance. Software often fails to take advantage of the full capabilities of the new format version.

Here, we describe Aird--A new format with controllable precision and information. It uses controllable precision and multiple index strategies to reconstruct the whole mass spectrum file, which makes it a more suitable data format for the computational process. The advantages of the Aird are as follows:

1. Faster decompression speed. Aird can decompress an m/z array of 400 megabytes per second;
2. Higher compression ratio, which can significantly reduce memory usage and bandwidth;
3. More flexible file reading strategies by multiple indexing strategies, which can improve block read speed and reduce memory usage;

Aird includes a metadata file and a spectrum data file. The metadata file contains the necessary information from the controllable vocabularies, which is stored as JSON file. For the large spectrum data (mainly m/z and intensity), Aird stores it with controllable precision. Users can set the necessary precision when converting the vendor files. After determining the required precision, Aird converts the floating-point m/z array to integer array. Because the m/z array is ordered in each spectrum, it is an effective way to compress relatively small deltas of adjacent integers instead of the large integers themselves. Aird uses the FastPfor library for differential coding of integer arrays (Lemire, Boytsov et al. 2016). The FastPfor library uses the Single Instruction Multiple Data (SIMD) technology, benefiting Aird to encode and decode the data faster with a high compression ratio.

Here we provide two tools for developers to use and understand Aird.

1. Aird SDK: The SDK of Aird data accessing for both Java and C# languages.
2. Aird Pro: A GUI client for data conversion from vendor file to Aird file. The vendor file reader API is from MSConvert(Adusumilli and Mallick 2017).

## Materials and Methods

An overview of the Aird data structure is shown in Fig.1 and the main methods are described here:

**Fig. 1.**
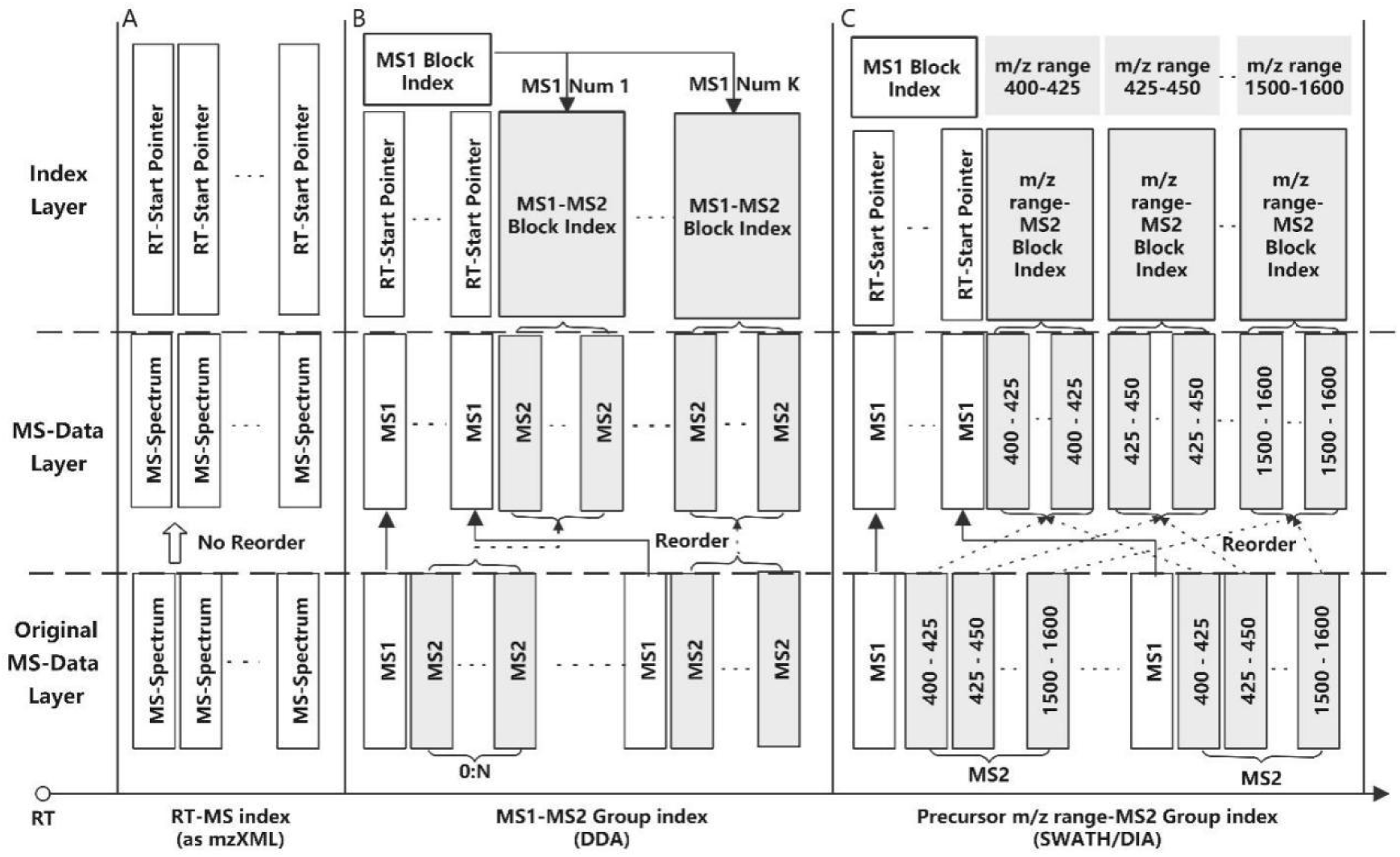
(A) Traditional mzXML-like indexing, MS data is sorted by retention time. The retention time and spectrum start position are stored as an index. (B) Generally used in DDA, each MS1 spectrum is followed by 0-N MS2 spectrums. Extracted ion chromatogram is generally calculated in successive MS1 spectrums, so reorder MS1 spectrums to put them in the same physical file block can speed up file reads. (C) Generally used in SWATH/DIA, every MS2 spectrum belongs to a certain m/z SWATH window. MS2 spectrums in the same window will be frequently accessed and calculated. Reordering is essential for fast file I/O.

### Multiple indexing and storage strategies

The storage of mass spectrometry data includes two parts: the index strategy and the actual storage location in the disk. For example, mzXML provides an index strategy by storing the scanning number and start position of each spectrum, as it stores specific MS data in the order of scanning number. Each scanning number represents a specific retention time (RT). The indexing strategy of mzXML is inflexible. Considering the Data Independent Acquisition (DIA) and Data Dependent Acquisition (DDA), Aird described three indexing and storage strategies:

1. RT-MS index: This is one of the most traditional indexing strategies, which is also used in mzXML and mzML format. This strategy is an efficient way to access successive spectrums. MS data are also linearly arranged by RT and stored in the file.
2. MS1-MS2 Group index: Considering DDA, each MS1 spectrum will correspond to zero or multiple MS2 spectrums. Adjacent MS1 spectrums are often taken out together to calculate the Extracted Ion Chromatogram (XIC). Therefore, Aird will reorder the spectrum data and put the data of MS1 together as one MS1 group. The MS2 groups corresponding to each MS1 will be put together. The start position of the MS1 group and MS2 group will be added as the new index data.
3. Precursor m/z range-MS2 Group index: Considering DIA/SWATH, each MS2 spectrum corresponds to a specified precursor m/z range. Aird also builds special index structures to speed up block reading for a particular m/z range. The MS2 groups corresponding to each specified precursor m/z range.

### JSON format for metadata storage

JSON is a lightweight data exchange format with similar readability and extensibility as XML. However, with the same semantics, JSON uses much less text than XML, which also makes the JSON file smaller (See Fig.2A). JSON is a subset of JavaScript, making it better in terms of web performance and parsing speed. Just like the mzXML Schema, we also provide a JSON Aird Schema(https://aird.oss-cn-beijing.aliyuncs.com/AirdMetaData.json). A detail description for every field in the Aird metadata is attached in the Supplementary Table.S1-Table.S8.

**Fig. 2.**
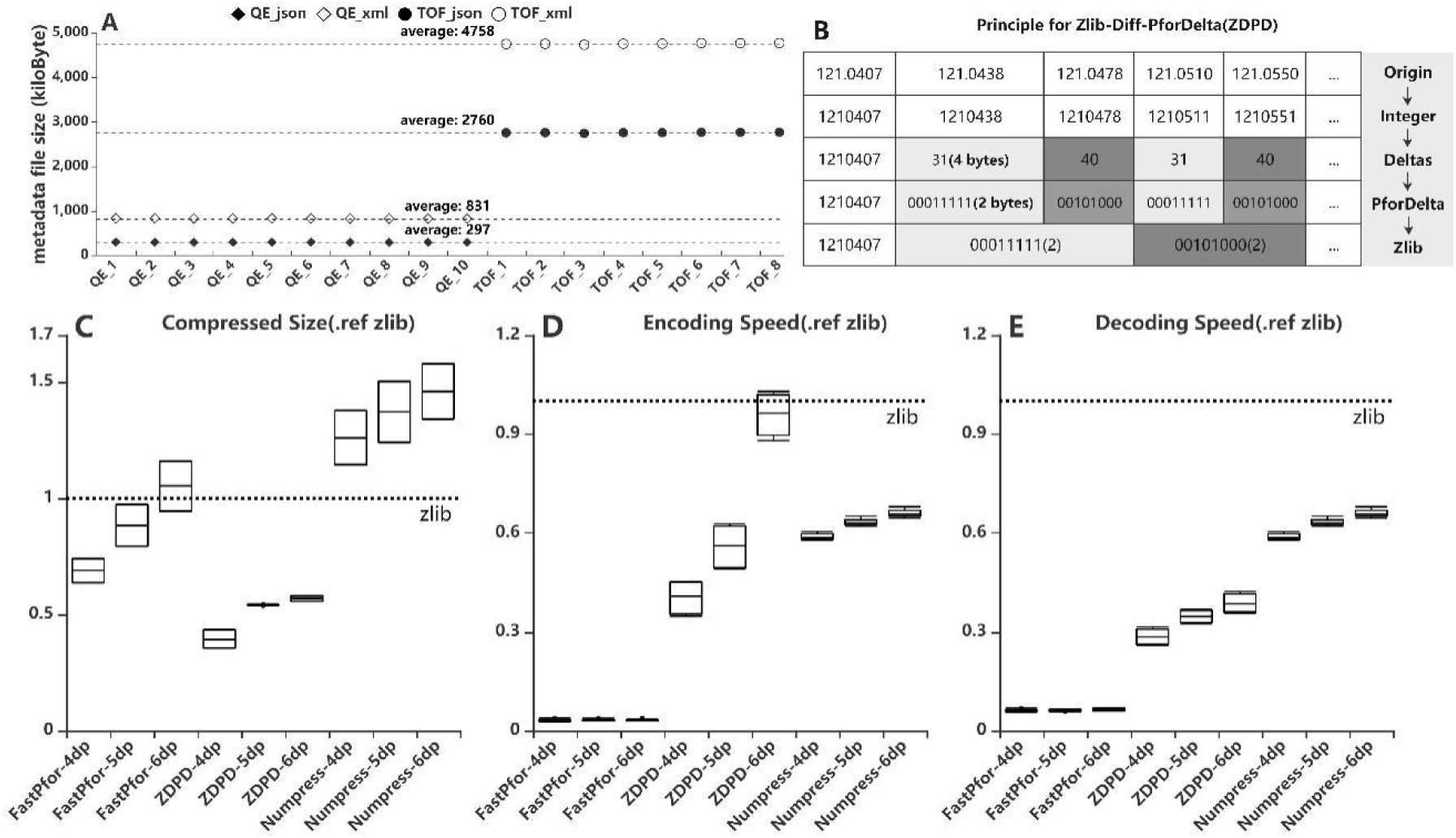
(A) Meta data with the same content is stored in format of json and xml, and the files generated by Thermo QE and Sciex TOF. File size increased by about 65% when transferred from json to xml for TOF files, and 1.7 times for files from Thermo QE. (B)The principle of ZDPD algorithm. (C) The compression size compared with Zlib using each compression algorithm. The ratio increases as the accuracy improves. ZDPD algorithm shows the best performance. (D) The encoding time for each algorithm. Due to SIMD support, the FastPfor algorithm is extremely fast. (E) The decoding time for each algorithm. Decoding time is one of the most important parameters in the computation - oriented process. Although ZDPD is not the fastest, the performance of ZDPD algorithm is the most balanced when combining the compression rate and decoding speed rate.

### Zlib-Diff-PforDelta (ZDPD) algorithm for m/z compression

The m/z array has the following characteristics

1. In metabolomics and proteomics, the fragment mass is generally between 0 and 3000;
2. The m/z arrays are ordered;
3. In m/z array, there will be several consecutive Numbers close to each other, and the difference of these Numbers is relatively small and fixed;

We build a new compressor ZDPD. The brief principle of ZDPD is shown in Fig.2B. Aird takes the integer type as the storage type for m/z array. If the needed data precision is 10 ppm (100 daltons to calculate), then the precision can be accurate to 3 decimal places (dp). If 0.1 ppm is needed, 5 dp is necessary. The m/z array in each spectrum is an ordered array. Aird achieves the best compression rate by combining FastPfor and Zlib algorithms reasonably. FastPfor library is a suitable compressor to store deltas between adjacent integers, benefiting from its special optimization. It also supports SIMD acceleration (Lemire, Boytsov et al. 2016). After compressing the sorted integers, Aird uses the Zlib algorithm to compress the delta values, which can achieve a better result than only using the Zlib or FastPfor algorithm. The combination of the two algorithms perfectly utilizes the prior characteristics of m/z array.

We test the data on an open dataset (Li, Lu et al. 2018) with a normal desktop computer (CPU: i7 7700K 4.4GHz, Disk: 5900R HDD, Memory:16GB). 8 files from AB SCIEX TripleTOF 6600 and 8 files from Thermo QE Orbitrap are mixed together for the test. The standard golden set (Rost, Rosenberger et al. 2014) and HYE datasets (Navarro, Kuharev et al. 2016) are also tested in the supplementary. The MSConvert version is 3.0.20196-20896b6b1

## RESULTS

Some open datasets (Li, Lu et al. 2018) are chosen to test the Aird algorithm. We compare data size, comparison ratio, encoding time, and decoding time with four different precision. Zlib, FastPfor, Aird, and Numpress are also compared (See Fig.2C,2D,2E).

Metadata information is usually read into the memory for quick preview of file information. Unlike mass spectrometry data, metadata usually exists in memory for a long time, while mass spectrometry data should be loaded into memory only during calculation. Although metadata is less than 5M, when loading a project contains hundreds of experimental files. It will also take up more than 500MB of memory. The JSON format can make file preview faster and reduce memory usage. XML and JSON format are used to store the same content to compare the storage size of the two formats (see Fig.2A). Due to the elimination of redundant tag information in XML, JSON format file is obviously smaller than XML format file.

In terms of actual size, the m/z data size compressed by the Aird algorithm is 65% of it of Zlib. When using OpenSWATH(Rost, Rosenberger et al. 2014), MZMine (Pluskal, Castillo et al. 2010), XCMS (Smith, Want et al. 2006) or other software, XIC is a very frequent and common calculation step. Due to the increasing size of MS data, it becomes more and more difficult to decode the mass spectrum data and put it into memory. Decoding speed has become one of the bottlenecks of the whole workflow. ZDPD algorithm also performs well in decoding speed due to the SIMD support. The decoding time is only 33% of it of Zlib. That means, with the same I/O strategy, Aird uses only half of the memory to complete the calculation compared with Zlib, but can increases the decompression speed by three times. For the intensity array, Aird uses the Zlib compressor with 1dp precision. As an option, Aird also offers an optional compression algorithm that can result in precision loss of up to 0.25%.

## DISCUSSION

### Comparison Ratio for Compressor

Digital properties of intensity arrays vary across instruments. Duplicate intensities are more tend to be produced by Time of Flight (TOF) spectrometer. This is a very favorable characteristic of the Zlib algorithm. Besides, m/z arrays with more numbers within the same range will have a higher compression rate as their delta values are smaller. Therefore, under the same conditions, the more the number of fragments contained in the sample, the higher the compression rate. So we can’t come up with a stable and absolute conclusion to express the exact compression performance of Aird files. It is also unfair to compare the absolute size between near-lossless and lossless formats. Although we provide some results regarding size comparison with mzXML and vendor files in the supplementary, they are only for readers’ reference.

### File format with supporting analysis platform

As we mentioned in the Introduction Chapter. Software often fails to take advantage of the full capabilities of the new format version. In addition to a compressor for m/z array, we also developed two omics platforms for Aird-Propro and MetaPro. Propro provides a whole workflow for DIA/SWATH and PRM proteomics, which is also a web-based platform with abundant visualization. Propro remains in internal testing, but you can follow the development progress in the Github (https://github.com/Propro-Studio/Propro) as it is an open-source project; MetaPro provides a whole process for targeted and untargeted metabolomics. By using Aird, both of the two platforms perform well in data processing.

Recent advances in proteome informatics have led to an explosion in tools to analyze mass spectrometry data. These tools operate across the analysis pipeline doing everything from assessing quality control to matching peptides to spectra to quantification. Unfortunately, the vast majority of these tools are not able to operate directly on the proprietary formats generated by the diverse mass spectrometers. Consequently, the first step in many protocols is the conversion of data from vendor-specific binary files to open-format files. This protocol details the use of ProteoWizard’s msConvert and msConvertGUI software for this conversion, taking format features, coding options, and vendor particularities into account. We specifically describe the various options available when doing conversions and the implications of each option.

The analysis and management of MS data, especially those generated by data independent MS acquisition, exemplified by SWATH-MS, pose significant challenges for proteomics bioinformatics. The large size and vast amount of information inherent to these data sets need to be properly structured to enable an efficient and straightforward extraction of the signals used to identify specific target peptides. Standard XML based formats are not well suited to large MS data files, for example, those generated by SWATH-MS, and compromise high-throughput data processing and storing. We developed mzDB, an efficient file format for large MS data sets. It relies on the SQLite software library and consists of a standardized and portable server-less single-file database. An optimized 3D indexing approach is adopted, where the LC-MS coordinates (retention time and m/z), along with the precursor m/z for SWATH-MS data, are used to query the database for data extraction. In comparison with XML formats, mzDB saves approximately 25% of storage space and improves access times by a factor of twofold up to even 2000-fold, depending on the particular data access. Similarly, mzDB shows also slightly to significantly lower access times in comparison with other formats like mz5. Both C++ and Java implementations, converting raw or XML formats to mzDB and providing access methods, will be released under permissive license. mzDB can be easily accessed by the SQLite C library and its drivers for all major languages, and browsed with existing dedicated GUIs. The mzDB described here can boost existing mass spectrometry data analysis pipelines, offering unprecedented performance in terms of efficiency, portability, compactness, and flexibility.

Summary Sorted lists of integers are commonly used in inverted indexes and database systems. They are often compressed in memory. We can use the single-instruction, multiple data (SIMD) instructions available in common processors to boost the speed of integer compression schemes. Our S4-BP128-D4 scheme uses as little as 0.7 CPU cycles per decoded 32-bit integer while still providing state-of-the-art compression. However, if the subsequent processing of the integers is slow, the effort spent on optimizing decompression speed can be wasted. To show that it does not have to be so, we (1) vectorize and optimize the intersection of posting lists; (2) introduce the SIMD GALLOPING algorithm. We exploit the fact that one SIMD instruction can compare four pairs of 32-bit integers at once. We experiment with two Text REtrieval Conference (TREC) text collections, GOV2 and ClueWeb09 (category B), using logs from the TREC million-query track. We show that using only the SIMD instructions ubiquitous in all modern CPUs, our techniques for conjunctive queries can double the speed of a state-of-the-art approach. Copyright © 2015 John Wiley & Sons, Ltd.

Data analysis represents a key challenge for untargeted metabolomics studies and it commonly requires extensive processing of more than thousands of metabolite peaks included in raw high-resolution MS data. Although a number of software packages have been developed to facilitate untargeted data processing, they have not been comprehensively scrutinized in the capability of feature detection, quantification and marker selection using a well-defined benchmark sample set. In this study, we acquired a benchmark dataset from standard mixtures consisting of 1100 compounds with specified concentration ratios including 130 compounds with significant variation of concentrations. Five software evaluated here (MS-Dial, MZmine 2, XCMS, MarkerView, and Compound Discoverer) showed similar performance in detection of true features derived from compounds in the mixtures. However, significant differences between untargeted metabolomics software were observed in relative quantification of true features in the benchmark dataset. MZmine 2 outperformed the other software in terms of quantification accuracy and it reported the most true discriminating markers together with the fewest false markers. Furthermore, we assessed selection of discriminating markers by different software using both the benchmark dataset and a real-case metabolomics dataset to propose combined usage of two software for increasing confidence of biomarker identification. Our findings from comprehensive evaluation of untargeted metabolomics software would help guide future improvements of these widely used bioinformatics tools and enable users to properly interpret their metabolomics results.

Consistent and accurate quantification of proteins by mass spectrometry (MS)-based proteomics depends on the performance of instruments, acquisition methods and data analysis software. In collaboration with the software developers, we evaluated OpenSWATH, SWATH 2.0, Skyline, Spectronaut and DIA-Umpire, five of the most widely used software methods for processing data from sequential window acquisition of all theoretical fragment-ion spectra (SWATH)-MS, which uses data-independent acquisition (DIA) for label-free protein quantification. We analyzed high-complexity test data sets from hybrid proteome samples of defined quantitative composition acquired on two different MS instruments using different SWATH isolation-window setups. For consistent evaluation, we developed LFQbench, an R package, to calculate metrics of precision and accuracy in label-free quantitative MS and report the identification performance, robustness and specificity of each software tool. Our reference data sets enabled developers to improve their software tools. After optimization, all tools provided highly convergent identification and reliable quantification performance, underscoring their robustness for label-free quantitative proteomics.

A broad range of mass spectrometers are used in mass spectrometry (MS)-based proteomics research. Each type of instrument possesses a unique design, data system and performance specifications, resulting in strengths and weaknesses for different types of experiments. Unfortunately, the native binary data formats produced by each type of mass spectrometer also differ and are usually proprietary. The diverse, nontransparent nature of the data structure complicates the integration of new instruments into preexisting infrastructure, impedes the analysis, exchange, comparison and publication of results from different experiments and laboratories, and prevents the bioinformatics community from accessing data sets required for software development. Here, we introduce the ‘mzXML’ format, an open, generic XML (extensible markup language) representation of MS data. We have also developed an accompanying suite of supporting programs. We expect that this format will facilitate data management, interpretation and dissemination in proteomics research.

BACKGROUND: Mass spectrometry (MS) coupled with online separation methods is commonly applied for differential and quantitative profiling of biological samples in metabolomic as well as proteomic research. Such approaches are used for systems biology, functional genomics, and biomarker discovery, among others. An ongoing challenge of these molecular profiling approaches, however, is the development of better data processing methods. Here we introduce a new generation of a popular open-source data processing toolbox, MZmine 2. RESULTS: A key concept of the MZmine 2 software design is the strict separation of core functionality and data processing modules, with emphasis on easy usability and support for high-resolution spectra processing. Data processing modules take advantage of embedded visualization tools, allowing for immediate previews of parameter settings. Newly introduced functionality includes the identification of peaks using online databases, MSn data support, improved isotope pattern support, scatter plot visualization, and a new method for peak list alignment based on the random sample consensus (RANSAC) algorithm. The performance of the RANSAC alignment was evaluated using synthetic datasets as well as actual experimental data, and the results were compared to those obtained using other alignment algorithms. CONCLUSIONS: MZmine 2 is freely available under a GNU GPL license and can be obtained from the project website at: http://mzmine.sourceforge.net/. The current version of MZmine 2 is suitable for processing large batches of data and has been applied to both targeted and non-targeted metabolomic analyses.

Metabolite profiling in biomarker discovery, enzyme substrate assignment, drug activity/specificity determination, and basic metabolic research requires new data preprocessing approaches to correlate specific metabolites to their biological origin. Here we introduce an LC/MS-based data analysis approach, XCMS, which incorporates novel nonlinear retention time alignment, matched filtration, peak detection, and peak matching. Without using internal standards, the method dynamically identifies hundreds of endogenous metabolites for use as standards, calculating a nonlinear retention time correction profile for each sample. Following retention time correction, the relative metabolite ion intensities are directly compared to identify changes in specific endogenous metabolites, such as potential biomarkers. The software is demonstrated using data sets from a previously reported enzyme knockout study and a large-scale study of plasma samples. XCMS is freely available under an open-source license at http://metlin.scripps.edu/download/.

The open XML format mzML, used for representation of MS data, is pivotal for the development of platform-independent MS analysis software. Although conversion from vendor formats to mzML must take place on a platform on which the vendor libraries are available (i.e. Windows), once mzML files have been generated, they can be used on any platform. However, the mzML format has turned out to be less efficient than vendor formats. In many cases, the naive mzML representation is fourfold or even up to 18-fold larger compared with the original vendor file. In disk I/O limited setups, a larger data file also leads to longer processing times, which is a problem given the data production rates of modern mass spectrometers. In an attempt to reduce this problem, we here present a family of numerical compression algorithms called MS-Numpress, intended for efficient compression of MS data. To facilitate ease of adoption, the algorithms target the binary data in the mzML standard, and support in main proteomics tools is already available. Using a test set of 10 representative MS data files we demonstrate typical file size decreases of 90% when combined with traditional compression, as well as read time decreases of up to 50%. It is envisaged that these improvements will be beneficial for data handling within the MS community.

The closed nature of vendor file formats in mass spectrometry is a significant barrier to progress in developing robust bioinformatics software. In response, the community has developed the open mzML format, implemented in XML and based on controlled vocabularies. Widely adopted, mzML is an important step forward; however, it suffers from two challenges that are particularly apparent as the field moves to high-throughput proteomics: large increase in file size, and a largely sequential I/O access pattern. Described here is ‘toffee’, an open, random I/O format backed by HDF5, with lossless compression that gives file sizes similar to the original vendor format and can be reconverted back to mzML without penalty. It is shown that mzML and toffee are equivalent when processing data using OpenSWATH algorithms, in additional to novel applications that are enabled by new data access patterns. For instance, a peptide-centric deep-learning pipeline for peptide identification is proposed. Documentation and examples are available at https://toffee.readthedocs.io, and all code is MIT licensed at https://bitbucket.org/cmriprocan/toffee.

Across a host of MS-driven-omics fields, researchers witness the acquisition of ever increasing amounts of high throughput MS data and face the need for their compact yet efficiently accessible storage. Addressing the need for an open data exchange format, the Proteomics Standards Initiative and the Seattle Proteome Center at the Institute for Systems Biology independently developed the mzData and mzXML formats, respectively. In a subsequent joint effort, they defined an ontology and associated controlled vocabulary that specifies the contents of MS data files, implemented as the newer mzML format. All three formats are based on XML and are thus not particularly efficient in either storage space requirements or read/write speed. This contribution introduces mz5, a complete reimplementation of the mzML ontology that is based on the efficient, industrial strength storage backend HDF5. Compared with the current mzML standard, this strategy yields an average file size reduction to approximately 54% and increases linear read and write speeds approximately 3-4-fold. The format is implemented as part of the ProteoWizard project and is available under a permissive Apache license. Additional information and download links are available from http://software.steenlab.org/mz5.

BACKGROUND: Mass Spectrometry (MS) is a widely used technique in biology research, and has become key in proteomics and metabolomics analyses. As a result, the amount of MS data has significantly increased in recent years. For example, the MS repository MassIVE contains more than 123TB of data. Somehow surprisingly, these data are stored uncompressed, hence incurring a significant storage cost. Efficient representation of these data is therefore paramount to lessen the burden of storage and facilitate its dissemination. RESULTS: We present MassComp, a lossless compressor optimized for the numerical (m/z)-intensity pairs that account for most of the MS data. We tested MassComp on several MS data and show that it delivers on average a 46% reduction on the size of the numerical data, and up to 89%. These results correspond to an average improvement of more than 27% when compared to the general compressor gzip and of 40% when compared to the state-of-the-art numerical compressor FPC. When tested on entire files retrieved from the MassIVE repository, MassComp achieves on average a 59% size reduction. MassComp is written in C++ and freely available at https://github.com/iochoa/MassComp. CONCLUSIONS: The compression performance of MassComp demonstrates its potential to significantly reduce the footprint of MS data, and shows the benefits of designing specialized compression algorithms tailored to MS data. MassComp is an addition to the family of omics compression algorithms designed to lessen the storage burden and facilitate the exchange and dissemination of omics data.

