## Supplemental File for "Aird: A computation-oriented mass spectrometry data format enables higher compression ratio and less decoding time"

### SUMMARY

#### Total File Size Comparison

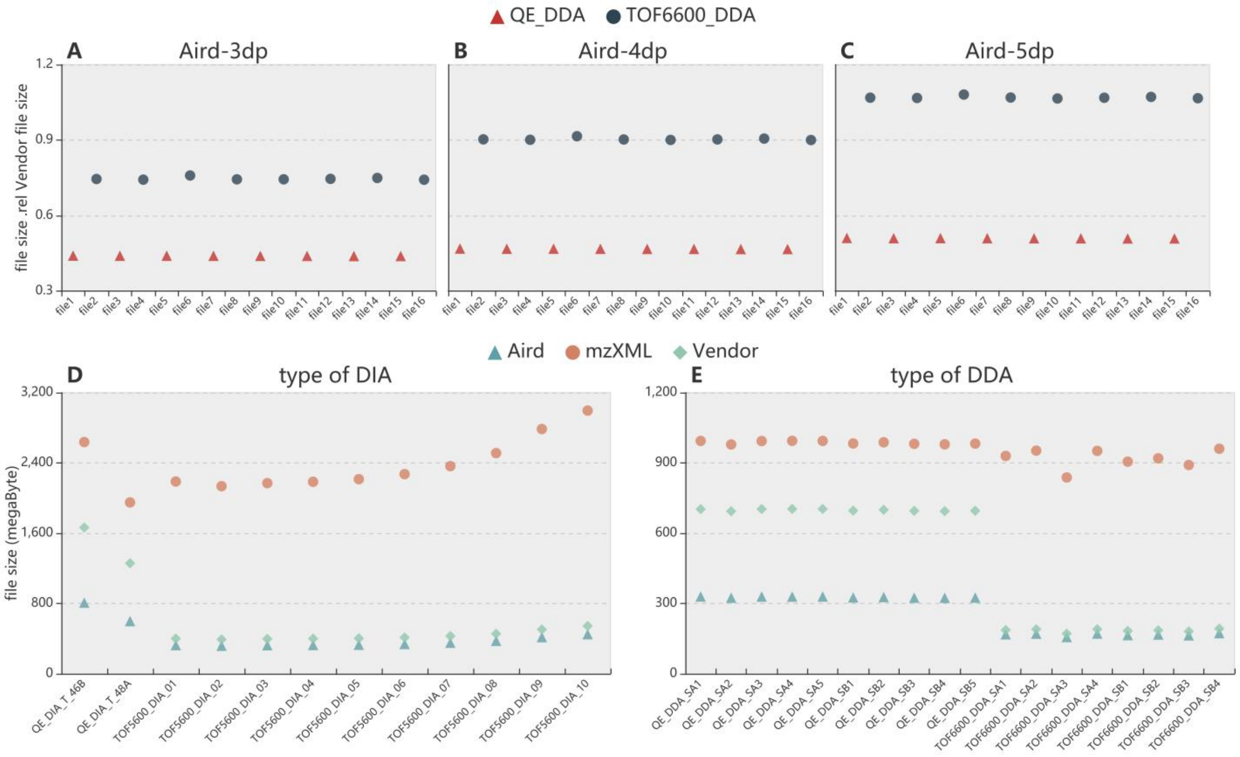


Fig.S1 Compressed file size is divided by its of Vendor file for Aird of different precision for Thermo QE and TOF DDA files(see in Fig.S1A, Fig.S1B and Fig.S1C). File size increases as the accuracy improves. Compared with 3 dp Aird files, 4 dp and 5 dp of QE data increased by 6.3% and 15.9% respectively in average. The corresponding increases for TOF6600 are 21.1% and 43.2%. Also, Aird of 4dp is compared with mzXML and vendor file for DIA and DDA files generated by Thermo QE and TOF( see in Fig.S1D and Fig.S1E). It's worth noting that file size of Aird is about 10% less than vendor file for data generated by TOF, and 50% when it comes to Thermo QE. In contrast, mzXML increased by 50% and 4 times.

### AirdPro

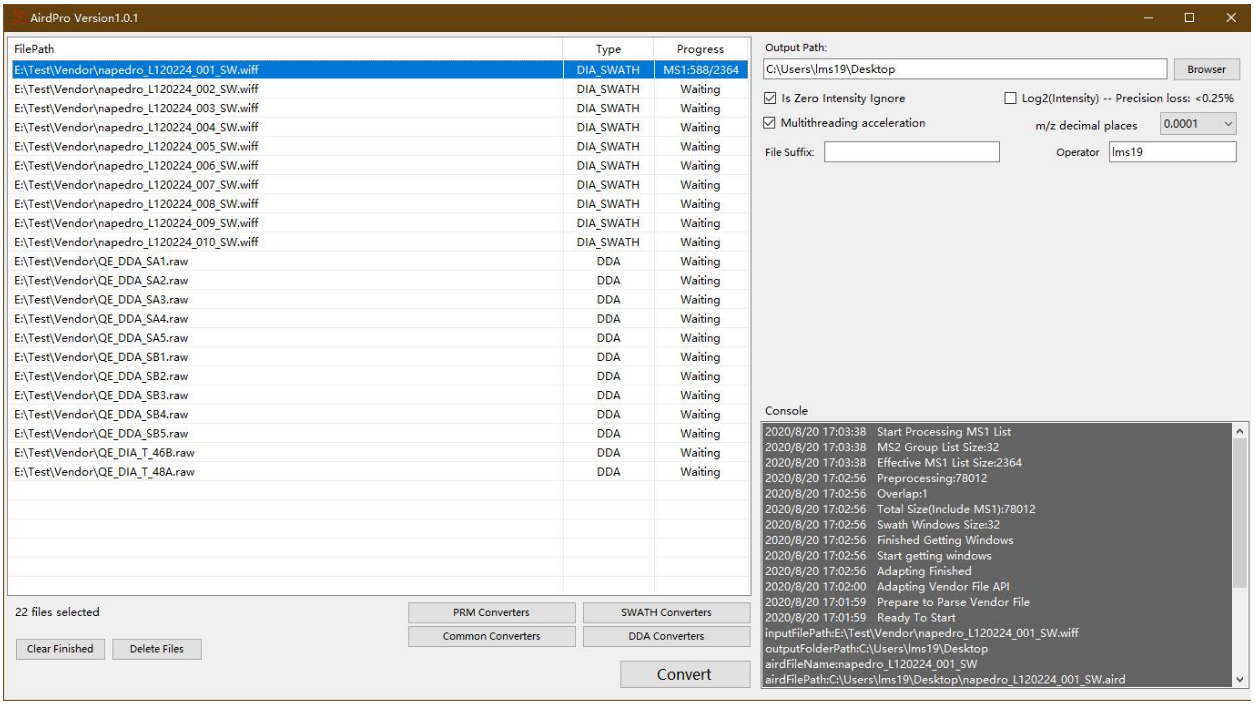


Fig.S2 The main form of AirdPro. Users can choose from four different file conversion modes:DDA, DIA/SWATH, PRM and Common mode.

AirdPro is written in the C# language, and the read capability for vendor files is based on pwiz_bindings_cli from the Proteowizard team. There are four action options on the right panel:

1. Is Zero Intensity Ignore: Choose whether to ignore values with an intensity of 0;
2. Log2(Intensity): If this option is used for precision approximation, it will bring about 0.25% precision loss.
3. Multithreading acceleration: Whether to use multithreading for conversion acceleration, which will have a significant acceleration on multi-core computers, but greatly improve CPU utilization.
4. m/z decimal places: Controllable precision option. Users can choose the exact number of digits for m/z data.

The log of the data conversion process can be viewed on the console.AirdPro can run on the Windows 7 or 10 with .NET Framework 4.7.2 or higher. Users can download the application from **https://aird.oss-cn-beijing.aliyuncs.com/AirdPro-V1.0.1.zip**.
AirdPro is opensource software which is under the Mulan PSL v2 (http://license.coscl.org.cn/MulanPSL2)

#### AirdSDK

AirdSDK is a Jar package for accessing all the information from the Aird file.AirdSDK is written in Java, which is a cross-platform language. The meta-structure of Aird is also in the package “bean”, The detail for the meta-structure is described here:

| Name | Type | Required | Description |
|---|---|---|---|
| compressors | List<Compressor> | True | The compression strategies for m/z and intensity array |
| rangeList | List<WindowRange> | False | The precursor m/z window ranges which have been adjusted with experiment overlap. This field is targeted for DIA and PRM type format |
| indexList | List<BlockIndex> | True | The index for mass spectrometry data |
| instruments | List<Instrument> | True | General information about the MS instrument |
| dataProcessings | List<DataProcessing> | False | Description of any manipulation (from the first conversion to Aird format until the creation of the current Aird instance document) applied to the data |
| softwares | List<Software> | False | Software used to convert the data. If data has been processed (e.g. profile > centroid) by any additional progs these should be added too |
| parentFiles | List<ParentFile> | False | Path to all the ancestor files (up to the native acquisition file) used to generate the current Aird document |
| version | String | True | Aird format version |
| versionCode | Integer | True | Aird format version code |
| type | String | True | Aird Type. There are four types now: DIA, DDA, PRM, COMMON |
| fileSize | Long | True | The file size for Aird file and JSON file |
| totalScanCount | Long | True | Total spectrums count |
| airdPath | String | False | The .aird file path |
| creator | String | False | The file creator, this field can be set up in the AirdPro |
| createDate | String | False | The create date for the aird file |
| ignoreZeroIntensityPoint | Boolean | True | Whether ignore the point which intensity is 0 |
| features | String | False | Some other features stored with "key:value;key:value" format |

Table.S1 AirdInfo Table

| Name | Type | Required | Description |
|---|---|---|---|
| target | String | True | mz, intensity |
| methods | List<String> | True | zlib, pFor, log10 |
| precision | Integer | False | 10^N, the N means N decimal places for the final data |
| byteOrder | String | True | LITTLE_ENDIAN, BIG_ENDIAN |

Table.S2 Compressor Table

| Name | Type | Required | Description |
|---|---|---|---|
| start | Double | True | Precursor m/z start |
| end | Double | True | Precursor m/z end |
| mz | Double | True | Precursor m/z |
| features | String | False | Some other features stored with “key:value;key:value” format |

Table.S3 WindowRange Table

| Name | Type | Required | Description |
|---|---|---|---|
| level | Integer | True | 1:MS1, 2:MS2 |
| startPtr | Long | True | The start point for the block |
| endPtr | Long | True | The endpoint for the block |
| num | Integer | False | The scan number in the vendor file. If a block has a list of MS2, this field is the related MS1’s number |
| rangeList | List<WindowRange> | False | The precursor m/z window ranges which have been adjusted with experiment overlap. This field is targeted for DIA and PRM type format |
| nums | List<Integer> | False | Scan numbers in the block |
| rts | List<Float> | True | All the retention times in the block |
| mzs | List<Long> | True | COMMON: the start position for every m/z bytes<br>Others: the size for every m/z bytes size |
| ints | List<Long> | True | COMMON: the start position for every m/z bytes<br>Others: the size for every m/z bytes size |
| features | String | False | Some other features stored with “key:value;key:value” format |

Table.S4 BlockIndex Table

| Name | Type | Required | Description |
|---|---|---|---|
| manufacturer | String | False | Instrument manufacturer: ”ABSciex”,”Thermo Fisher” |
| ionization | String | False | Ionization |
| resolution | String | False | Resolution |
| model | String | False | Instrument model |
| source | List<String> | False | Source: ”electrospray ionization”, ”electrospray inlet” |
| analyzer | List<String> | False | Analyzer: “quadrupole”, “orbitrap” |
| detector | List<String> | False | Detector: ”inductive detector” |
| features | String | False | Some other features stored with “key:value;key:value” format |

Table.S5 Instrument Table

| Name | Type | Required | Description |
|---|---|---|---|
| processingOperations | List<String> | False | Any additional manipulation not included elsewhere in the dataProcessing element |

Table.S6 DataProcessing Table

| Name | Type | Required | Description |
|---|---|---|---|
| name | String | True | The software name |
| version | String | False | The software version |

Table.S7 Software Table

| Name | Type | Required | Description |
|---|---|---|---|
| name | String | True | The filename |
| location | String | False | The file location |
| type | String | False | The file type |

Table.S8 ParentFile Table

We also provide a JSON schema file of Aird index file for developers:
**https://aird.oss-cn-beijing.aliyuncs.com/AirdMetaData.json**
The Aird SDK jar package can be download:
**https://aird.oss-cn-beijing.aliyuncs.com/aird-sdk.jar**

### Aird File Conversion Workflow

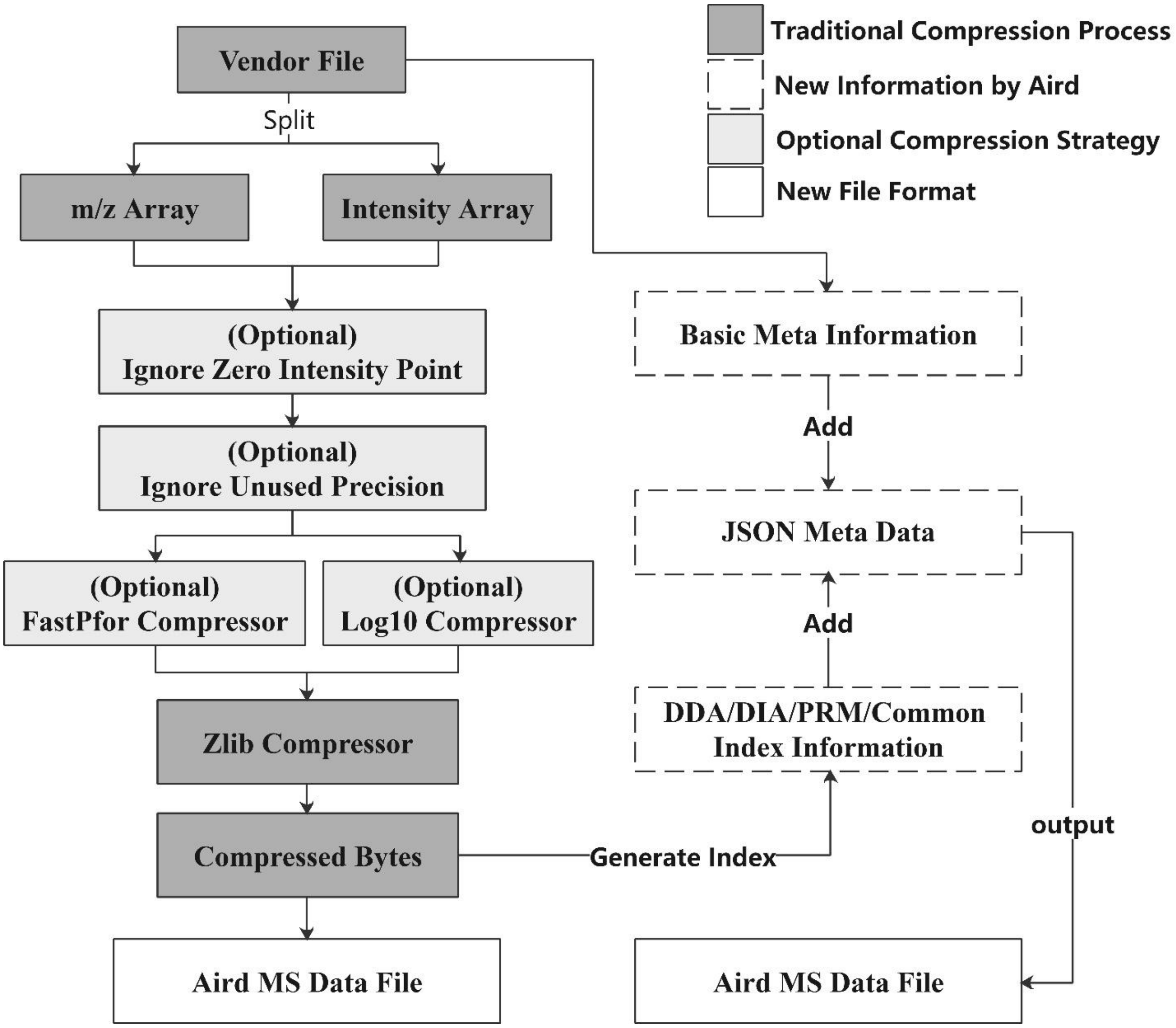


Fig.S3 The conversion workflow for AirdPro. The picture shows how AirdPro separated the vendor files and compressed the m/z array and intensity array.

### Framework

#### *AirdPro*

| Framework | Version | License | How to get |
|---|---|---|---|
| .NET | 4.7.2 | Free to use | https://dotnet.microsoft.com/download/dotnet-framework |
| Json | 12.0.1 | MIT | https://www.newtonsoft.com/json |
| Zlib | 1.1.3 | GNU LGPL | https://archive.codeplex.com/?p=dotnetzip |
| FastPfor | 0.1.12 | Apache V2.0 | https://github.com/Genbox/CSharpFastPFOR |

Table.S9 Main third party libraries used in AirdPro

Except for C#, AirdPro uses 3 third-party libraries(see Table.S9). AirdSDK is written by Java. Except for Java, AirdSDK uses 3 thrid-party libraries(see Table.S10).

#### *AirdSDK*

| Framework | Version | License | How to get |
|---|---|---|---|
| OpenJDK | 14 | GPL V2 | https://jdk.java.net/14/ |
| FastPfor | 0.1.12 | Apache V2.0 | https://github.com/lemire/JavaFastPFOR |
| Fastjson | 1.2.56 | Apache V2.0 | https://github.com/alibaba/fastjson |
| Common-lang3 | 3.7 | Apache V2.0 | http://commons.apache.org/proper/commons-lang/ |

Table.S10 Main third party libraries used in AirdSDK

### Datasets

In this paper, parts of the following public datasets are used. The data files are shown in Table.S11.

| Description | License<br>How to get |
|---|---|
| SWATH-MS Gold Standard Dataset | https://www.peptideatlas.org/PASS/PASS00289 |
| Hybrid proteome samples of Human, Yeast, E.coli | https://www.ncbi.nlm.nih.gov/pubmed/27701404 |
| DDA samples from two platforms:<br>1. AB SCIEX TripleTOF 6600 interfaced with Shimazu L30A UPLC<br>2. Thermo Q Exactive HF with Dionex UltiMate 3000 HPLC | https://drive.google.com/drive/folders/1PRDIvihGFgkmErp2fWe41UR2Qs2VY_5G?usp=sharing_eip&ts=5b8ab35f<br>(TripleTOF 6600 dataset ID: 1197236, QE HF dataset ID: 1197351) |

Table.S11 Datasets compared in the paper
